# The Tsallis index of the human cortical transcriptome is invariant between bipolar II disorder and control: a pre-registered, cross-platform null with the disease question relocated to correlation structure

**DOI:** 10.64898/2026.07.21.739779

**Authors:** Sergio Assunção Monteiro, Fabricio Alves Barbosa da Silva

## Abstract

Gene-expression fluctuations are heavy-tailed and resist description by additive (Boltzmann–Gibbs) statistics, motivating a non-extensive (Tsallis) account in which the index *q* summarizes the departure from Gaussianity. We ask two separable questions of the human cortical transcriptome in bipolar II disorder: whether the model-free value of *q* differs between cases and controls, and, if not, whether any disease signal instead resides in the correlation structure. Using a per-sample maximum-likelihood *q*-Gaussian estimator under a leakage-free, pre-registered protocol, we find the index concentrated near 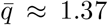 (GSE80655), 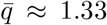 (GSE12649), 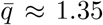 (GSE53987), with no case–control difference in any cohort. Per-sample model comparison favours the *q*-Gaussian over a Gaussian in 99–100% of samples (median ΔBIC ≪ 0), so *q* is a meaningful descriptor rather than an artefact of fitting. The BD–CTRL difference is null in every cohort. Pooling the three homogeneous cohorts, we test invariance by equivalence (TOST) rather than by non-rejection: the data establish equivalence at the bound |Δ*q*| ≥ 0.037, but do *not* reach the pre-registered bound of 0.03, which would require roughly twice the present sample. We report this shortfall explicitly: the null is bounded and informative, but the study is underpowered against its own pre-registered minimum effect of interest. A random-matrix test finds no between-group structural difference surviving label permutation. We argue that non-extensivity behaves as a conserved organizational property of the cortical transcriptome and that the disease question, for bipolar II, is relocated from the marginal index to the collective correlation modes and to dynamics (companion work). We report, and do not paper over, a failed cross-tissue scale anchor: a glioma RNA-seq reference is depth-confounded in a cohort-inconsistent way and therefore cannot license a same-value claim across tissues.

## 1 Introduction

Non-extensive statistical mechanics generalizes the Boltzmann–Gibbs framework through the Tsallis entropy and its associated index *q* [1]; the *q*-Gaussian arises as the maximum-*S*_*q*_ distribution under a variance constraint and as an attractor in a *q*-generalized central limit theorem for suitably correlated variables [4, 3], making *q* a natural single-number summary of heavy-tailed fluctuations [2, 5].

Prior work on tumor (glioblastoma) transcriptomes reported that *q* is a robust, cohort-independent property of the data recoverable model-free, with the disease signature expressed in the reorganization of gene–gene correlations rather than in the marginal index. That observation motivates, but does not fix the value of, the present study: we ask whether the same organizing principle holds in *non-tumor* human cortex, and whether the signature of bipolar II disorder lives in *q* or in correlation geometry.

We treat the two questions as pre-registered hypotheses. H1: the per-sample *q* differs between bipolar and control prefrontal cortex. H2: the gene–gene correlation structure differs between groups. The design fixed, before any confirmatory analysis, the estimator, the decision rule, the acceptable null, and the giving-up criterion; a two-branch outcome was specified in advance (a conserved-constant branch and a biomarker branch). *Pre-registered threshold, honestly reported:* the preregistration fixed a minimum effect of interest of |Δ*q*| *<* 0.03. We retain that threshold as the target and assess it with a formal equivalence test (TOST) rather than with non-rejection of the null, which would only be absence of evidence. As reported below, the pooled data do not reach the pre-registered bound; we say so rather than substituting a weaker criterion after seeing the result. We follow a convention-free rule throughout: because the absolute value of *q* depends on platform and pre-processing, we never compare absolute *q* across platforms, and we re-establish any inherited value independently within the same pipeline rather than citing it.

## 2 Methods

### Cohorts and stratum

We analyze three independent cortical cohorts spanning two platforms: two microarray series – GSE12649 (prefrontal, Brodmann 46) [10] and GSE53987 (restricted to the prefrontal stratum) [11] – and one RNA-seq series, GSE80655 (dorsolateral prefrontal cortex) [12]. All analyses are restricted to the prefrontal stratum for cross-cohort comparability; per-cohort *n* is reported after quality control (below), and differs from the deposited totals because of stratum restriction and QC. Two glioma RNA-seq cohorts (CGGA 325/693) [13, 14] are processed by the identical pipeline solely as a cross-tissue scale reference; as reported in Results they fail QC and are excluded from the confirmatory family.

### Estimator

For each sample, deviations from the cohort-median expression profile are fitted to a *q*-Gaussian, 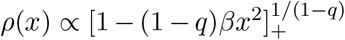, by maximum likelihood over (*q, β*). Maximum likelihood is used rather than a moment estimator because the *q*-Gaussian kurtosis diverges at *q* = 7/5, within the observed range. The scale parameter *β* and the joint identifiability of (*q, β*) (Hessian conditioning, multi-start agreement) are reported in the Supplementary material.

### Pre-processing (convention-free)

RNA-seq counts are filtered to expressed genes (counts ≥ 10 in ≥ 50% of samples; cf. [8, 9]), then transformed to log_2_(CPM + 1); microarray intensities are used as provided. We do *not* apply rarefaction in the main pipeline: because CPM already normalizes for sequencing depth, rarefaction-to-minimum is redundant and, over a wide depth range, distorts the estimate by discarding information unevenly; we verified (Supplementary) that the invariance conclusion is unchanged under log-CPM alone, rarefaction-to-minimum, and rarefaction-to-median. This is a within-platform measurement: absolute *q* is never compared across platforms.

### Model justification (H0 for the framework)

Per sample, we compare the *q*-Gaussian against a Gaussian by AIC and BIC (the *q*-Gaussian carries one extra parameter and is thus penalized) and report the Gaussian Kolmogorov–Smirnov statistic as a descriptive goodness-of-fit. This tests whether a non-extensive description is warranted at all rather than assumed.

### Technical confound

Per-sample 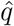 is tested against library size and zero-count fraction (bootstrap Spearman) ; the estimate is placed on the same processing scale as the confirmatory analysis, and both the analysis-scale and raw-scale correlations are reported (Supplementary QC figure) . A cohort whose 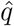 remains depth-confounded on the analysis scale is flagged and excluded from the confirmatory family (fail-fast).

### Structure (H2)

The gene –gene sample-covariance eigenspectrum is compared to the Marchenko–Pastur band 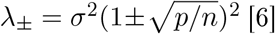 eigenvalues above *λ*_+_ are counted as collective modes, following applications of random-matrix theory to gene-expression covariance [19, 20]. Because *p* ≫ *n* and the data are heavy-tailed (so the fourth moment need not exist), the MP edge is used only as a reference: the between-group inference rests on a group-label permutation test of the mode count and of the leading eigenvalue, not on the absolute edge.

### Power

For each confirmatory cohort we report the minimum detectable effect (MDE) at 80% power, *α* = 0.05, both analytically (two-sample) and by re-running the actual permutation test under an injected shift; we report the more conservative (larger) value.

### Statistics

Effect sizes with bootstrap confidence intervals (*B* ≥ 5000), label-permutation nulls (10^4^ for H1; group-label permutation with fixed per-test seeds for H2), and Benjamini–Hochberg FDR [7] across the pre-registered confirmatory family; analyses use NumPy and SciPy [17, 18]. All numbers in this manuscript are injected from a re-executable report; none are hand-typed.

## 3 Results

### A non-extensive description is warranted

Per sample, the *q*-Gaussian is favoured over a Gaussian by BIC in 99% of GSE80655 samples (median ΔBIC = −1688); 100% of GSE12649 samples (median ΔBIC = −1058); 100% of GSE53987 samples (median ΔBIC = −5231), despite the extra-parameter penalty.

### No BD–CTRL difference in the Tsallis index

In GSE80655 (*n* = 47: 23 BD, 24 CTRL), 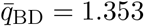 vs 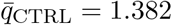 (Δ*q* = −0.030; Cohen’s *d* = −0.26, 95% CI [-0.90, 0.31]; permutation *p* = 0.371, FDR *q* = 0.592).

In GSE12649 (*n* = 67: 33 BD, 34 CTRL), 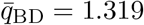 vs 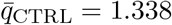 (Δ*q* = −0.019; Cohen’s *d* = −0.21, 95% CI [-0.70, 0.26]; permutation *p* = 0.395, FDR *q* = 0.592).

In GSE53987 (*n* = 36: 17 BD, 19 CTRL),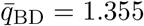 vs 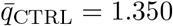 (Δ*q* = 0.005; Cohen’s *d* = 0.08, 95% CI [-0.60, 0.81]; permutation *p* = 0.806, FDR *q* = 0.806).

### Pooled effect and formal equivalence

Because the three cohorts are statistically homogeneous (*I*^2^ = 0%, *Q* = 0.70, *p* = 0.71), we pool them by inverse-variance meta-analysis of Hedges’ *g* (small-sample corrected) . The pooled effect is *g* = −0.153 (95% CI [-0.469, +0.163]; Δ*q* = −0.0136, 95% CI [-0.0417, +0.0145]; *p* = 0.34), and pooling lowers the minimum detectable effect from |Δ*q*| ≈ 0.059– 0.093 per cohort to |Δ*q*| = 0.0407 (Fig. 4). We test invariance by two one-sided tests (TOST), which is an equivalence test, rather than by non-rejection of the null, which is only absence of evidence. Against the pre-registered minimum effect of interest (|Δ*q*| *<* 0.03), TOST does *not* establish equivalence (*p* = 0.126): the study is underpowered relative to its own pre-registration. What the data do establish is equivalence at the bound |Δ*q*| ≥ 0.0371 (90% CI of the pooled effect) . Reaching the pre-registered bound of 0.03 would require approximately 159 subjects per group (available here: 73 BD / 77 CTRL). We state this explicitly rather than relaxing the criterion after the fact.

**Figure 1.**
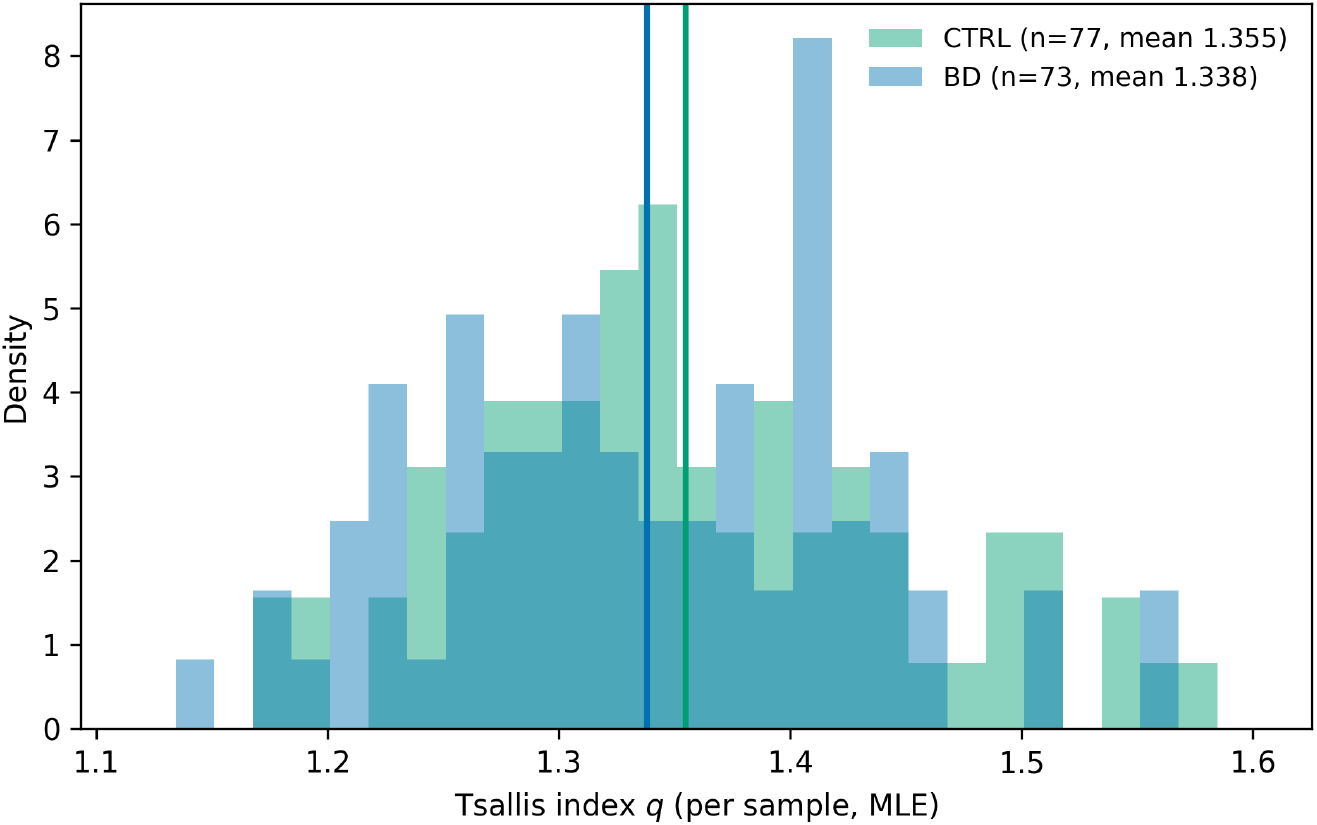
Per-sample Tsallis index *q* by group (maximum likelihood). Vertical lines mark group means.

**Figure 2.**
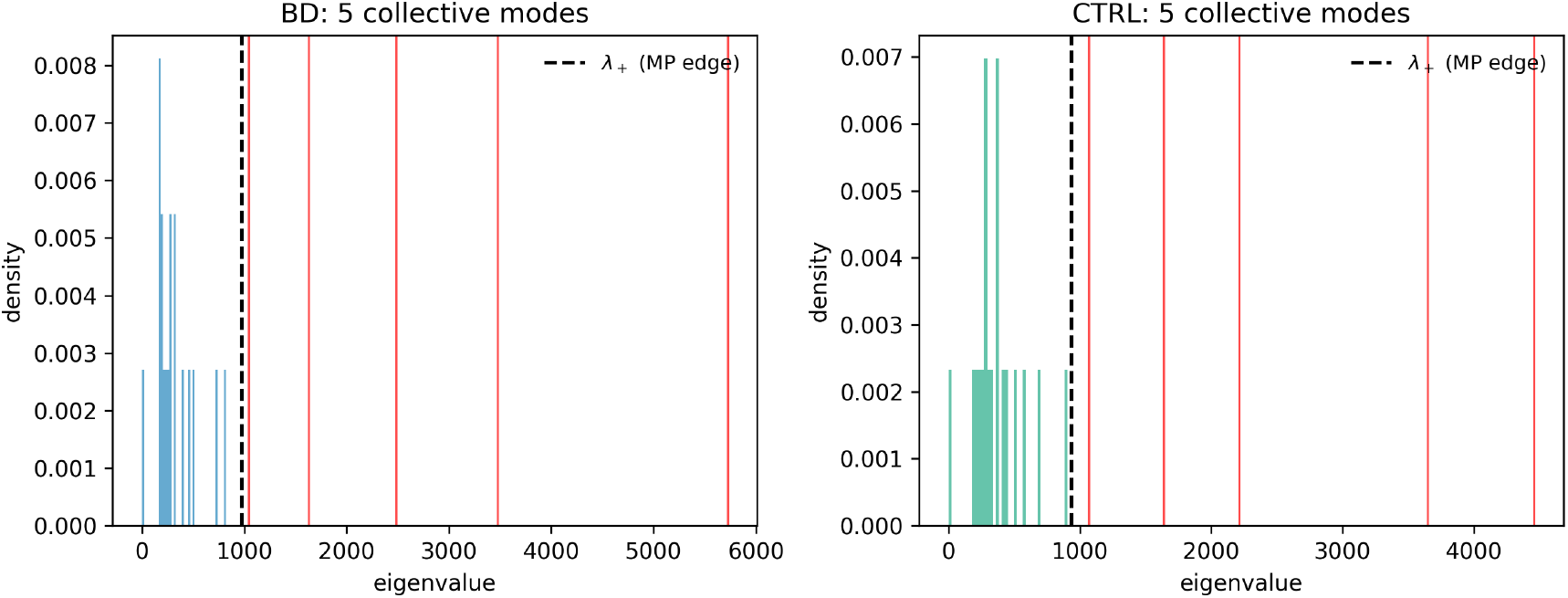
Gene–gene covariance eigenspectrum versus the Marchenko–Pastur reference band; dashed line marks *λ*_+_ and red lines mark eigenvalues above the edge. The edge is a reference only; inference uses label permutation (Methods).

### What the null does and does not exclude

Per cohort, the minimum detectable effect at 80% power is |Δ*q*| ≈ 0.061–0.135 (Cohen’s *d* between 0.68 and 0.94), which lies *above* the pre-registered minimum effect of interest (0.03): no single cohort could have detected the effect the study set out to find (Fig. 3). Pooling repairs part of this, but not all of it. The null is therefore bounded and quantitative, but it is bounded at 0.037, not at 0.03.

**Figure 3.**
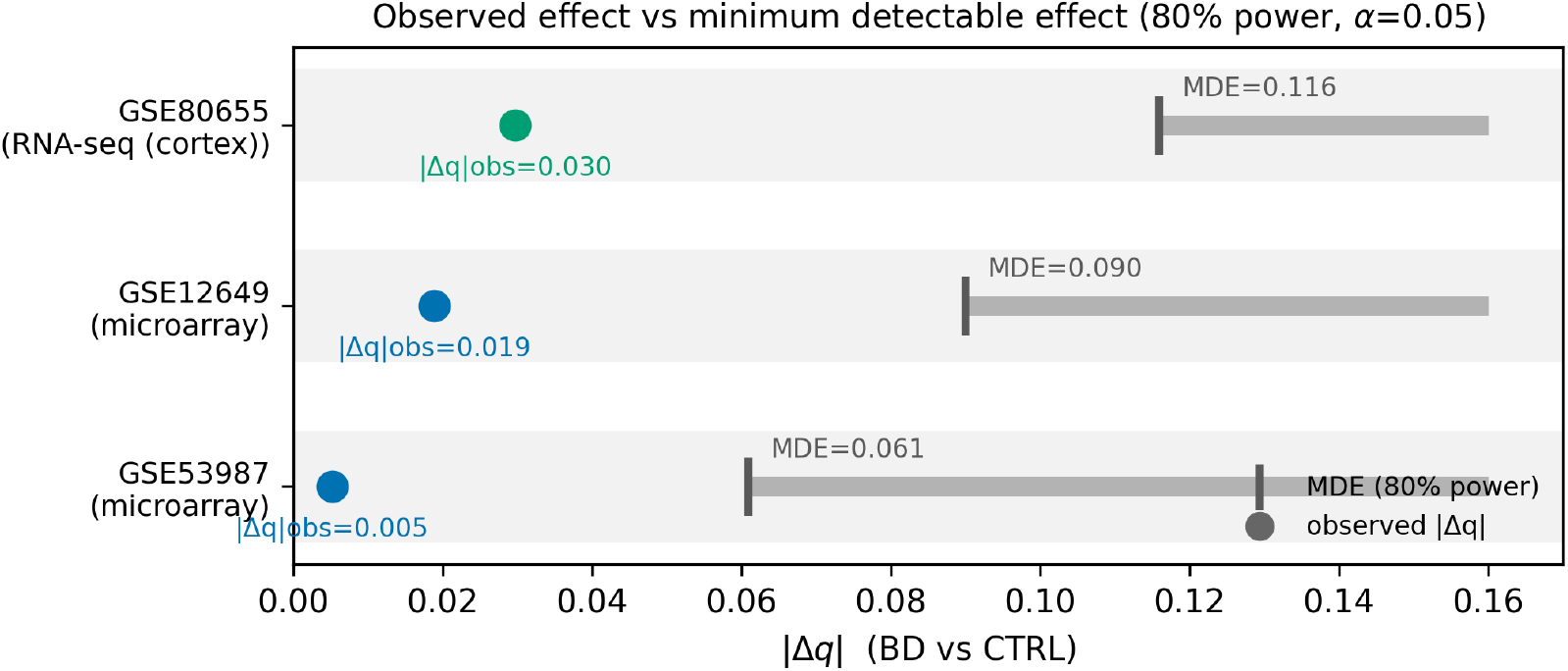
Observed |Δ*q*| (BD vs CTRL) versus the minimum detectable effect at 80% power (*α* = 0.05) per confirmatory cohort. Note that the per-cohort MDE exceeds the pre-registered minimum effect of interest (|Δ*q*| = 0.03); the pooled analysis (Fig. 4) is what bounds the null quantitatively.

**Figure 4.**
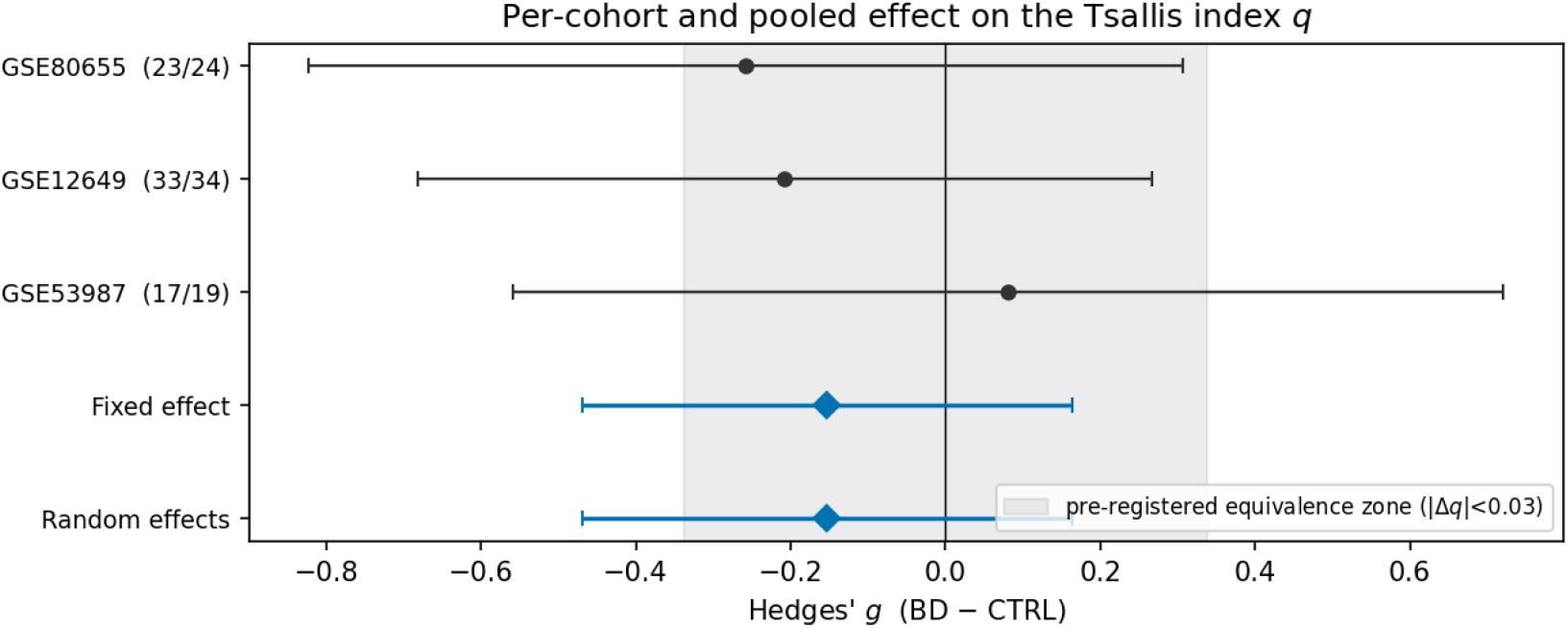
Per-cohort and pooled effect of diagnosis on the Tsallis index *q* (Hedges’ *g*, small-sample corrected; 95% CI). The shaded band is the pre-registered equivalence zone (|Δ*q*| *<* 0.03). Pooling is justified by the absence of heterogeneity (*I*^2^ = 0%). The pooled interval is not contained in the pre-registered zone: the null is bounded, but not at the pre-registered threshold.

### No structural signal

GSE80655: 5 vs 5 modes (*p*_modes_ = 1.000, *p*_*λ*_ = 0.173); GSE12649: 7 vs 8 modes (*p*_modes_ = 0.555, *p*_*λ*_ = 0.269); GSE53987: 5 vs 4 modes (*p*_modes_ = 0.422, *p*_*λ*_ = 0.724). No between-group difference in collective-mode count or leading eigenvalue survives label permutation in any cohort.

### A cross-tissue scale anchor fails, honestly

Run through the identical pipeline, glioma RNA-seq (CGGA) yields per-sample *q* in the same ∼1.3–1.4 window as cortex (Fig. 5); however 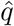 is confounded by sequencing depth in a cohort-inconsistent way (CGGA325 *ρ* = 0.29 [0.13, 0.43]; CGGA693 *ρ* = 0.27 [0.15, 0.39]), and the partial correlation conditioning on tumor covariates (IDH, 1p/19q, age, histology) removes the confound in one cohort but not the other. We therefore do *not* use glioma as a scale anchor and make no same-value claim across tissues; the numerical proximity is reported as suggestive only, consistent with the convention-free rule.

**Figure 5.**
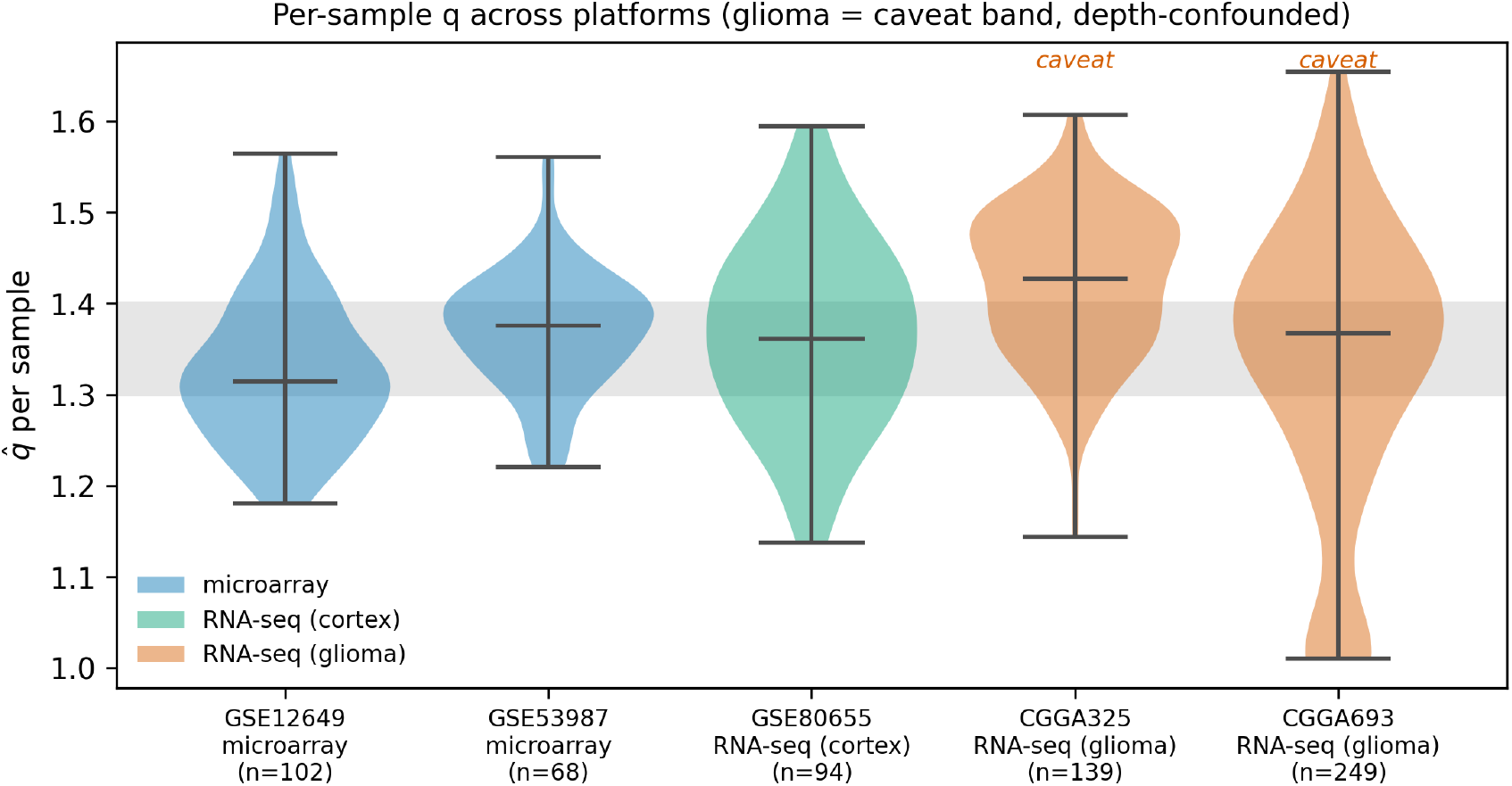
Per-sample *q* across cohorts and platforms. Cortical microarray and RNA-seq concentrate in the same ∼1.3–1.4 window (within-cortex, cross-platform invariance) . Glioma RNA-seq (orange) is shown as a caveat: its 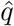 is depth-confounded and it is not used as a cross-tissue anchor.

**Figure 6.**
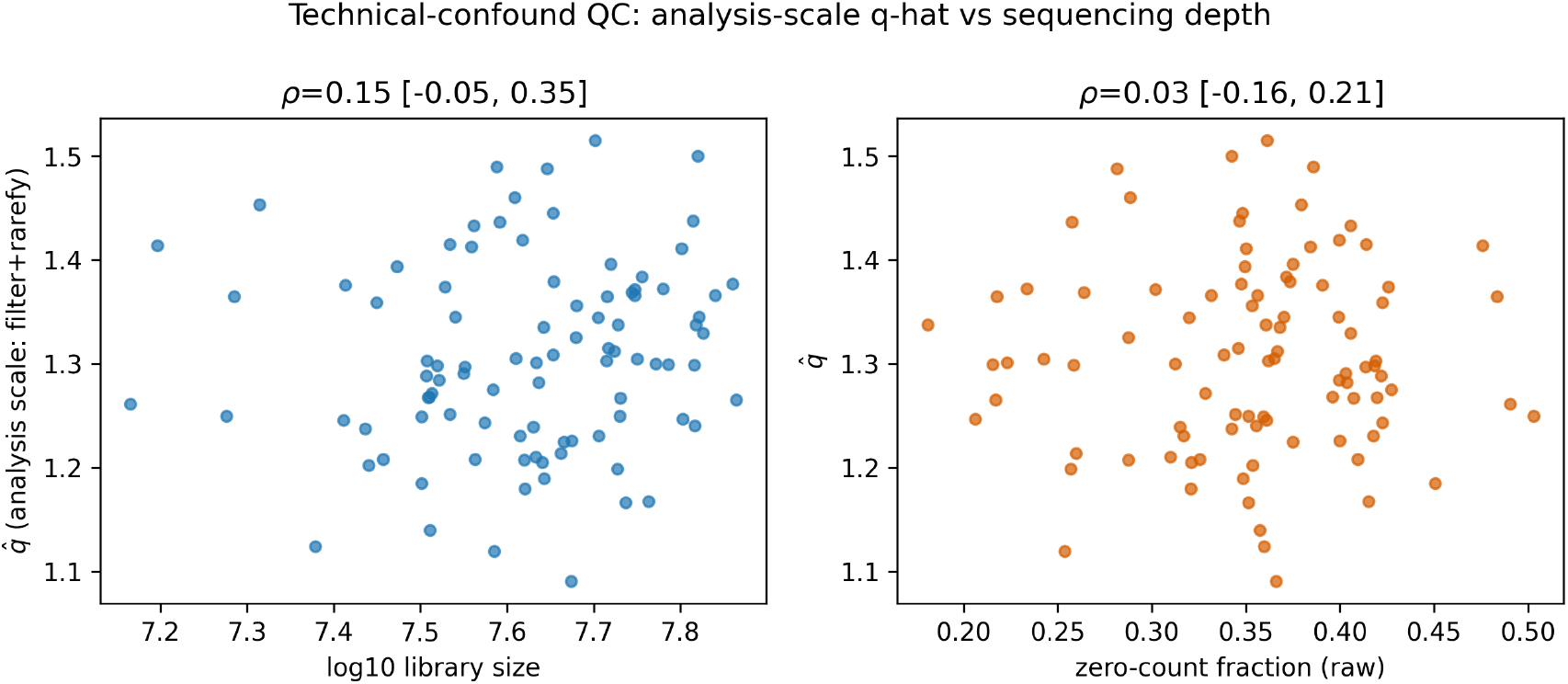
Technical-confound QC for the cortical RNA-seq cohort (GSE80655): per-sample 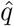 versus sequencing depth on the analysis scale (filter+CPM) and, for transparency, on the raw scale; the expression filter removes the raw-scale depth dependence. QC for the glioma cohorts (which remain depth-confounded) is shown in the Supplementary material.

## 4 Discussion

Under the pre-registered decision rule, the outcome is branch R1 (conserved property) : no detectable BD-vs-CTRL difference in q (Cohen’s d CI includes zero; |dq| well below the 80%-power MDE) in every confirmatory cohort, replicated across two platforms; no structural signal survives permutation – consistent with q behaving as a conserved organizational property of the cortical transcriptome. The result is a pre-registered null: across three independent cohorts and two platforms, the model-free Tsallis index of the prefrontal transcriptome does not differ between bipolar II and control, and no between-group structural signal survives permutation. We are deliberate about the strength of this statement. Equivalence is established at |Δ*q*| ≥ 0.037 but not at the pre-registered 0.03; effects in the interval between these bounds remain possible and are not excluded by these data.

Because the *q*-Gaussian is strongly preferred over the Gaussian per sample, this is a statement about a genuine non-extensive descriptor, not about a fitting artefact.

For bipolar II specifically, the contribution is one of principled elimination: for this family of marginal statistics, the cortical post-mortem transcriptome carries no detectable case–control signal, which relocates the search for a disease signature to the correlation geometry (collective modes) and to critical *dynamics*, the target of the companion EEG work [21, 22, 23, 24]; this is consistent with cross-disorder transcriptomic changes being expressed at the network level [15] and with the highly polygenic, distributed genetic architecture of bipolar disorder [16]. Read positively, an invariant *q* is a candidate organizational constant of the cortical transcriptome, providing a normality baseline against which individual, dynamic departures — rather than group means — may later be assessed.

A state-versus-trait boundary bounds this claim. The post-mortem transcriptome is a trait-level, structural snapshot taken at an unknown and presumably inter-episodic mood state; it cannot register a transient, state-dependent reorganization of expression during an active episode (for example a hypomanic switch), which could lower the dynamic phenotype’s complexity without moving the trait-level marginal index we measure. Our invariance is therefore a statement about the trait level, and is consistent with — not contradicted by — a state-dependent effect that only longitudinal, dynamic measurement can resolve. This is the same boundary that motivates relocating the disease question to dynamics and to cortical criticality accessible by EEG, where a non-extensive index behaves as an organizing quantity while group differences remain small and are expressed in the criticality relation rather than in the marginal value [21, 22, 23, 24]; read through the complexity-paradigm lens [25], an invariant trait-level *q* is what one expects of a conserved organizational property, with the pathological signature sought in dynamics.

### Limitations

(i) The random-matrix test is coarse and operates in a *p* ≫ *n*, heavy-tailed regime where the Marchenko–Pastur edge is only a reference; the structural null is limited to the two statistics tested under permutation and does not imply “no structure”. (ii) Absolute *q* is platform- and pipeline-dependent; our claim is within-cortex invariance across platforms, not a universal value, and explicitly not a cross-tissue same-value claim (the glioma anchor failed QC). (iii) The sample size is a ceiling of the public domain, not a design choice: post-mortem bipolar transcriptomes derive from few brain collections, and the widely reused series that appear to offer free reinforcement (GSE5388/5392; GSE92538) are re-analyses of the *same donors* as cohorts we already include (Stanley Array Collection and Pritzker, respectively) . Pooling them would be pseudo-replication rather than added power; we audited provenance before any aggregation (Supplementary). (iv) The study is underpowered against its own pre-registered minimum effect of interest: equivalence holds at |Δ*q*| ≥ 0.037 but not at 0.03, and closing that gap requires roughly twice the present number of donors. Because the per-sample estimator is precise (Supplementary), the limiting variance is between-subject and biological: only more donors, not deeper sequencing, can tighten this bound. (v) The positive control in the released pipeline is a synthetic surrogate of the estimator; the real-glioma control was run but is depth-confounded and therefore not a clean anchor. (vi) Post-mortem covariates (pH, PMI, RIN) are not fully modelled; a within-cohort sensitivity analysis is provided in the Supplementary material. (vii) The superstatistical reading of *q* is exploratory. (viii) The design is trait-level by construction: post-mortem tissue is a structural snapshot at an unknown mood state and cannot exclude a transient, state-dependent complexity change during an active episode; only longitudinal or dynamic measurement (companion EEG work) can address that regime.

## Supporting information

https://zenodo.org/records/21296799

## Acknowledgments

We thank the data-generating consortia and depositors whose openly available resources made this work possible: the submitters of the GEO series GSE12649, GSE53987 and GSE80655, and the Chinese Glioma Genome Atlas (CGGA). Computation was performed on local resources at PROCC/FIOCRUZ.

## CRediT authorship contribution statement

**Sergio Assunção Monteiro:** Conceptualization, Methodology, Software, Formal analysis, Investigation, Data curation, Visualization, Writing – original draft. **Fabricio Alves Barbosa da Silva:** Conceptualization, Supervision, Resources, Validation, Writing – review & editing.

## Data availability

All code, the per-sample derived indices, the locked pre-registration protocol, and a re-executable report that regenerates every number in the manuscript are openly available at Zenodo (DOI: 10.5281/zenodo.21296799; concept DOI, versioned on acceptance). Publicly available GEO series (GSE12649, GSE53987, GSE80655) are obtained from their providers and not redistributed here; the glioma cohorts (CGGA-325/693), used only as a diagnostic control, are governed by their consortium’s terms, with derived indices deposited rather than raw matrices. No controlled-access data are redistributed. The figures and tables reproduce offline from the deposited derived tables.

## Declaration of generative AI and AI-assisted technologies in the manuscript preparation process

During the preparation of this work the authors used Claude (Anthropic) to assist with language editing and drafting, with literature search and verification of bibliographic details, and with development of the analysis and figure-generation code. After using this tool the authors reviewed and edited the content as needed and take full responsibility for the content of the published article.

