## Supplementary material for "The Tsallis index of the human cortical transcriptome is invariant between bipolar II disorder and control: a pre-registered, cross-platform null with the disease question relocated to correlation structure": https://zenodo.org/records/21296799

Numbers and tables below are injected from re-executable artefacts by `code/build_supplement.py`; none are hand-typed. Generated: `2026-07-10T19:17:12`.

#### Contents

**1 Per-sample $q$ by diagnostic group (all diagnoses)** **1**

**2 Identifiability of the $(q, \beta)$ estimate** **1**

**3 Glioma (CGGA) as a rejected cross-tissue scale anchor** **2**

**4 Robustness to normalization choice** **3**

#### 1 Per-sample $q$ by diagnostic group (all diagnoses)

Figure 1 in the main text restricts to the pre-registered contrast (bipolar II vs control). For completeness, Fig. S1 shows the per-sample maximum-likelihood Tsallis index $q$ for every diagnosis available in the pooled cohorts (CTRL, BD, and the off-contrast SCZ/MDD groups where present). The off-contrast groups are shown for reference only and are not part of any confirmatory test.

#### 2 Identifiability of the $(q, \beta)$ estimate

The per-sample estimator fits $(q, \beta)$ jointly by maximum likelihood. Identifiability of $q$ rests on two properties, shown in Fig. S2: (A) the profile log-likelihood in $q$—with $\beta$ profiled out—has a single, well-separated maximum (sharp curvature, not a flat plateau); and (B) the per-sample standard error $\mathrm{se}(\hat{q})$ is small. We report the $q$–$\beta$ correlation (C) only as context: a *tilted* confidence ellipse ($|r|$ away from a degenerate value) is still identifiable provided the profile is peaked, so the correlation is not itself the test.

In GSE80655 ($n = 281$), the per-sample $q$ is estimated with median $\mathrm{se}(\hat{q}) = 0.009$ and 100% of fits converged; the profile likelihood peaks sharply in $q$ (panel A), so $q$ is identifiable. The $q$–$\beta$

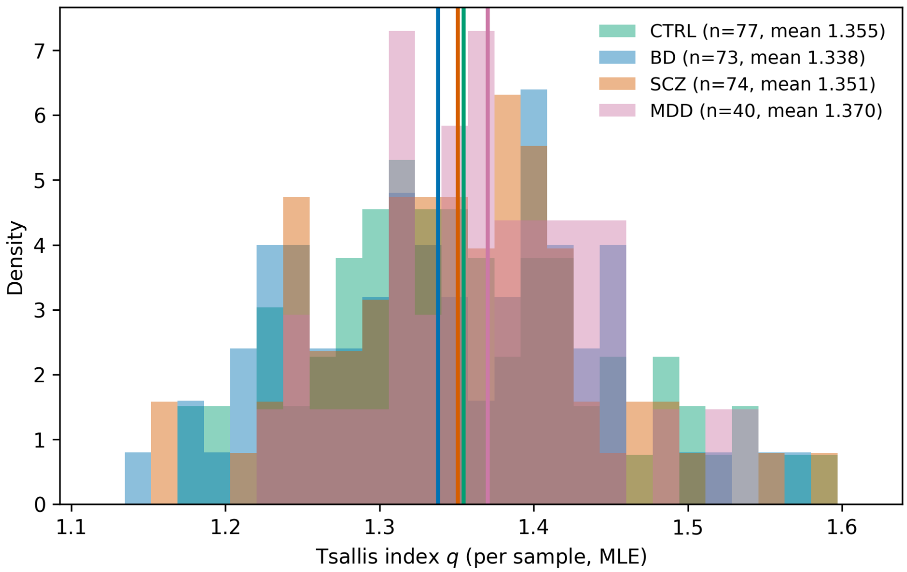


Figure S1: Per-sample $q$ across all diagnostic groups (maximum likelihood). BD and CTRL are the pre-registered contrast; SCZ/MDD are off-contrast and shown for reference.

confidence region is tilted but non-degenerate (median $|r| = 0.80$, max 0.86): a correlated, not a flat, ridge.

*Scope.* This diagnostic uses the full GSE80655 series (all regions), as identifiability of the estimator is region-agnostic; the confirmatory analysis in the main text is restricted to the DLPFC stratum.

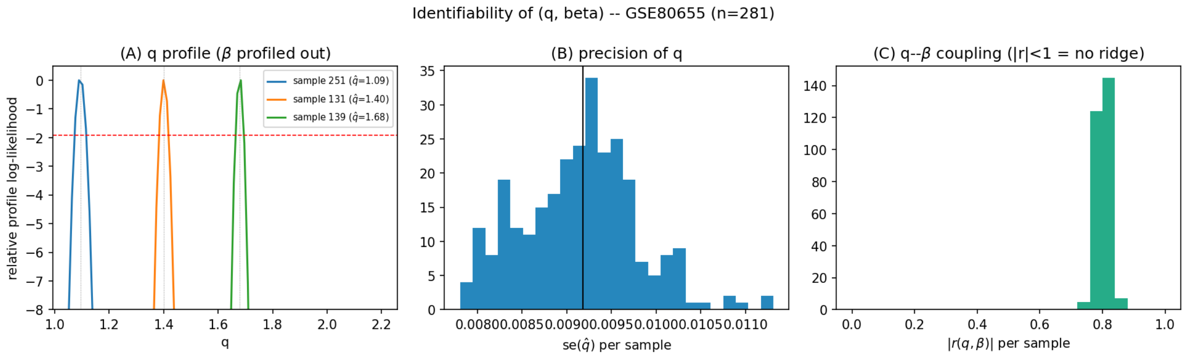


Figure S2: Identifiability of $(q, \beta)$. (A) Per-sample profile log-likelihood in $q$ ($\beta$ profiled out) for representative low/median/high-$q$ samples; dashed line marks a ~95% drop. (B) Distribution of $\mathrm{se}(\hat{q})$. (C) Distribution of the $q$–$\beta$ correlation $|r|$ (context, not the identifiability test).

#### 3 Glioma (CGGA) as a rejected cross-tissue scale anchor

We attempted to anchor a cross-tissue $q$-scale comparison using glioma RNA-seq (CGGA 325 and CGGA 693), processed through the *identical* cortical pipeline. Although the per-sample $q$ values fall in the same numerical window as cortex, the glioma estimate is confounded by sequencing depth, and—critically—the confound behaves *inconsistently across the two cohorts* once clinical covariates are partialled out. We therefore do not use glioma as an anchor and make no same-value claim across tissues.

Table S1 reports, per cohort, the raw Spearman correlation $\rho(\hat{q}, \log\text{-depth})$ on the analysis scale and the partial correlation conditioning on IDH mutation status, 1p/19q codeletion, age, histology, and sex (linear regression + Spearman of residuals). In CGGA 693 the confound is removed by conditioning (partial CI includes zero), consistent with tumour biology (purity/CNV) rather than a purely technical effect; in CGGA 325 it persists (partial CI excludes zero). This disagreement is the reason glioma cannot serve as a clean scale reference.

| cohort | $n$ | $\rho_{\text{raw}}$ | 95% CI | $\rho_{\text{partial}}$ | 95% CI | confound persists |
|---|---|---|---|---|---|---|
| CGGA325 | 139 | +0.309 | [+0.154, +0.452] | +0.323 | [+0.169, +0.466] | yes |
| CGGA693 | 249 | +0.265 | [+0.140, +0.384] | +0.054 | [−0.069, +0.185] | no |

Table S1: Depth confound of the glioma $\hat{q}$: raw vs. covariate-adjusted (partial) Spearman correlation with log-depth on the analysis scale. Partial correlation conditions on IDH, 1p/19q, age, histology, and sex. "Confound persists" is true when the partial-correlation 95% CI excludes zero.

Supporting the biological reading, the per-sample $q$ tracks IDH status: CGGA325 ($\tilde{q}$ by IDH: Mutant 1.409, Wildtype 1.447); CGGA693 ($\tilde{q}$ by IDH: Mutant 1.334, Wildtype 1.376). QC scatter plots of $\hat{q}$ versus sequencing depth for each glioma cohort are shown in Figs. S3–S4.

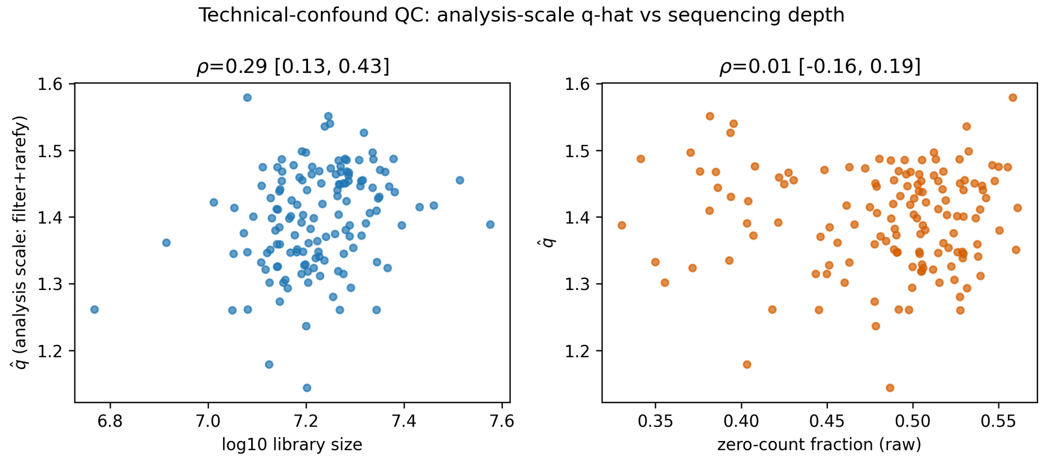


Figure S3: CGGA 325: per-sample $\hat{q}$ versus log-depth (analysis scale).

### 4 Robustness to normalization choice

The confirmatory pipeline uses $\log_2$(CPM+1) after low-expression filtering, with rarefaction removed from the default (CPM already normalizes; rarefaction is an opt-in flag). Fig. S5 shows that the per-sample $q$ and the depth correlation are stable across normalization variants (plain log-CPM, rarefy-to-minimum, and rarefy-to-median), so the reported invariance is not an artefact of the normalization choice. Table S2 gives the per-cohort median $q$ and depth correlation for each scheme. *Scope:* the depth diagnostic runs on the full GSE80655 series (all regions); the weak cortical correlation

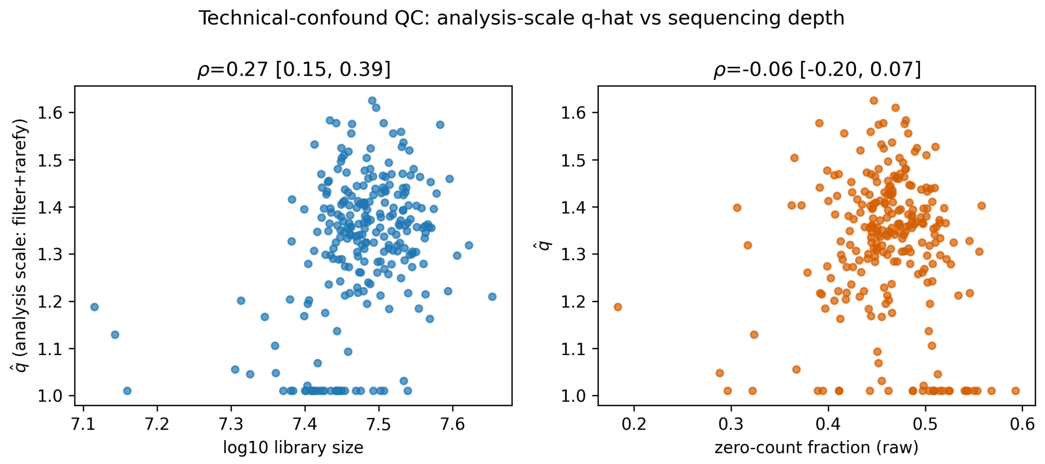


Figure S4: CGGA 693: per-sample $\hat{q}$ versus log-depth (analysis scale).

($\rho \approx 0.16$) is below the pre-registered confound threshold ($|\rho| > 0.2$) under every scheme, and the confirmatory DLPFC QC (main text) has a CI crossing zero.

| cohort | scheme | median $q$ | $\rho(\hat{q}, \log\text{-depth})$ | 95% CI | confound |
|---|---|---|---|---|---|
| CGGA325 | logCPM_only | 1.427 | +0.309 | [+0.154, +0.452] | yes |
| | rarefy_min | 1.403 | +0.291 | [+0.135, +0.434] | yes |
| | rarefy_median | 1.417 | +0.317 | [+0.163, +0.457] | yes |
| CGGA693 | logCPM_only | 1.367 | +0.265 | [+0.140, +0.384] | yes |
| | rarefy_min | 1.349 | +0.268 | [+0.145, +0.388] | yes |
| | rarefy_median | 1.359 | +0.268 | [+0.144, +0.387] | yes |
| GSE80655 | logCPM_only | 1.401 | +0.162 | [+0.050, +0.273] | no |
| | rarefy_min | 1.326 | +0.098 | [−0.018, +0.210] | no |
| | rarefy_median | 1.372 | +0.165 | [+0.051, +0.277] | no |

Table S2: Per-sample $q$ and its depth correlation under three normalization schemes. “Confound” is yes when $|\rho| > 0.2$ and the 95% CI excludes zero.

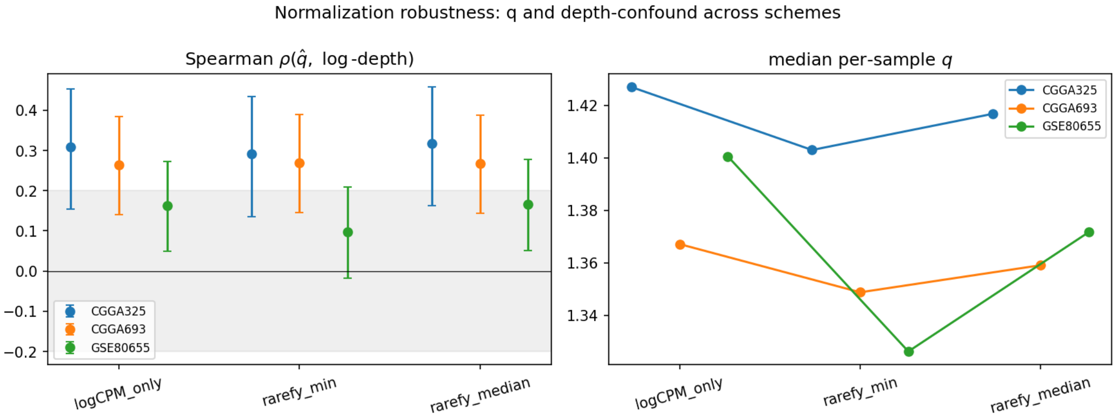


Figure S5: $q$ (right) and the depth correlation $\rho(\hat{q}, \log\text{-depth})$ with 95% CI (left) under three normalization schemes; the shaded band marks $|\rho| \leq 0.2$.
